# Anticodon-like loop-mediated dimerization in the crystal structures of HDV-like CPEB3 ribozymes

**DOI:** 10.1101/2022.09.22.508989

**Authors:** Anna Ilaria Przytula-Mally, Sylvain Engilberge, Silke Johannsen, Vincent Olieric, Benoît Masquida, Roland K.O. Sigel

**Affiliations:** Department of Chemistry, University of Zurich, CH-8057 Zurich, Switzerland; Swiss Light Source, Paul Scherrer Institute, CH-5232 Villigen, Switzerland; UMR 7156, CNRS - Université de Strasbourg, IPCB, 21 rue René Descartes, Strasbourg 67084, France

**Keywords:** RNA, HDV ribozyme, CPEB3 ribozyme, pseudoknot, dimerization

## Abstract

Cytoplasmic polyadenylation element-binding (CPEB) proteins are involved in many cellular processes, including cell division, synaptic plasticity, learning, and memory. A highly conserved, short mammalian ribozyme has been found within the second intron of the CPEB3 gene. Based on its cleavage mechanism and structural features, this ribozyme belongs to the hepatitis delta virus (HDV)-like ribozyme family. Here, we present the first crystallographic structures of human and chimpanzee CPEB3 ribozymes, both confirming the general topology of the HDV ribozyme with two parallel coaxial helical stacks. However, the residues involved in forming the P1.1 mini-helix, which is an integral part of the characteristic nested double pseudoknot involving P1, P2, and P3, instead participate in a seven nucleotides loop with a conformation similar to the one from the anticodon (AC) loop of tRNAs when interacting with the mRNA codon. The conformation of the loop supports the formation of a four-base pair helix by interacting with the AC-like loop from a symmetry-related ribozyme leading to ribozyme dimer formation. The present crystal structures link for the first time the sequence specificities of the CPEB3 and the HDV (genomic and antigenomic) ribozymes to their different structural features. This work corroborates the hypothesis made by Szostak that HDV ribozymes may have evolved from the CPEB3 ribozyme.

## INTRODUCTION

Since its discovery, the genomic HDV ribozyme has been one of the most studied model small self-cleaving ribozymes, contributing to a better understanding of the physicochemical basis of the self-scission mechanism, illustrating the principles of metal ion/nucleic acid interaction and more generally RNA structural features. The first crystal structure of the genomic HDV ribozyme, obtained from the post-cleavage state, revealed that it comprises five double-stranded A-helical regions (P1, P1.1, P2, P3, and P4) joined by three single-stranded regions (J1/2, J1.1/4, and J4/2) organized into two parallel helical stacks where P1/P1.1/P4 constitute one of them, and P2/P3 the other (Ferré-D’Amaré et al. 1998a). The structure also uncovered the prominent role of the P1.1 mini-helix and the P3-P1 cross-over in the organization of the catalytic core (**Figure 1A**). The highly complex nested double pseudoknot (P1/P2, P1.1/P3) weaves the two helical stacks together and builds up a platform on top of which sits the catalytic site, notably the catalytically important C75 (Perrotta and Been 1991; Ferré-D’Amaré et al. 1998a; Perrotta et al. 1999).

**Figure 1.**
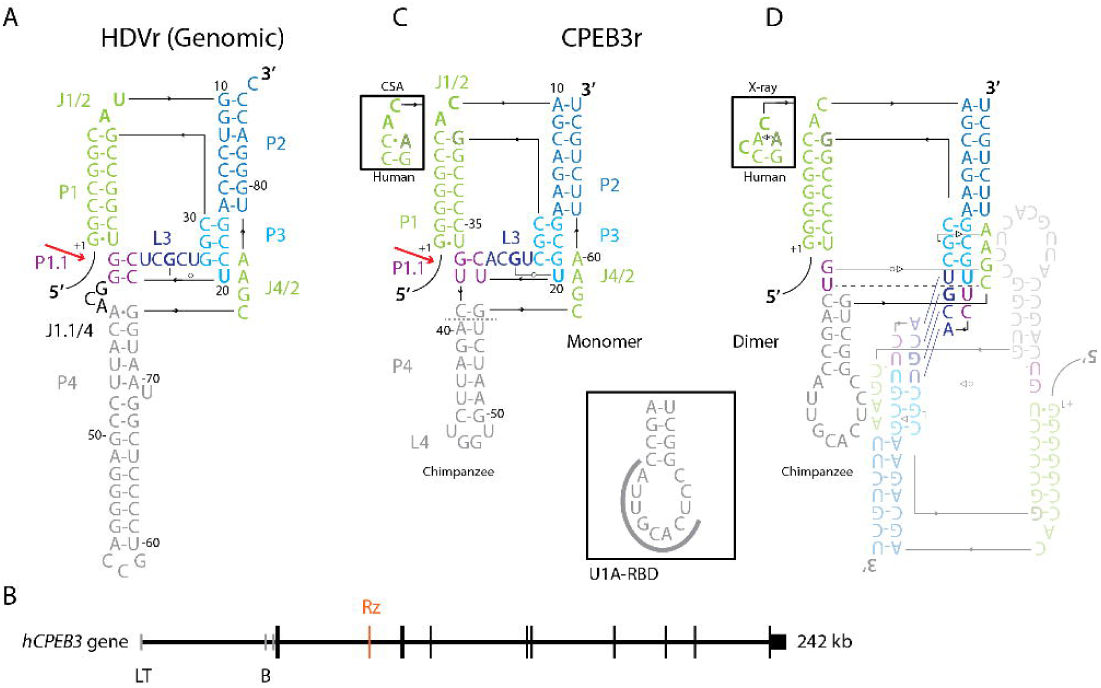
Secondary structure of (**A**) the genomic HDV ribozyme with individually colored secondary structure elements. The red arrow indicates the cleavage site. (**B**) Mapping of the human *CPEB3* gene. Horizontal lines (black) correspond to introns. Tissue specific exons are shown by gray vertical lines (L stands for liver, T for testis, and B for the brain). Translated exons are shown by black vertical lines (black). The CPEB3 ribozyme (Rz) location within the second intron is marked by an orange vertical line (adapted from Salehi-Ashtiani (Salehi-Ashtiani et al. 2006)). **C**) The secondary structure model of the CPEB3 wild-type ribozyme as deduced from the HDV ribozyme crystal structure suggests the presence of a weak P1.1 mini-helix containing a G-C pair and a putative U-U base pair deduced by comparison to the genomic HDV ribozyme crystal structure. The single nucleotide difference between human (A_30_) and chimpanzee (G_30_) CPEB3 ribozymes is shown in the upper inset, which shows the secondary structure of the human J1/2 region deduced from comparative sequence analysis (CSA). Non-canonical base pairs are highlighted using symbols from the Leontis-Westhof nomenclature (Leontis and Westhof 2001). The replacement of P4/L4 by the U1A-RBD used for crystallographic purposes is shown in the lower inset and takes place right below the dashed line shown on the WT model. The secondary structure model of the humanCPEB3 ribozyme J1/2 as deduced from the crystal structure is shown in the upper left corner inset. The connectivity between nucleotides is drawn using black arrows. The leader sequence preceding the cleavage site is represented by a black curve. P stands for base-paired regions, L for loops, and J for single-stranded joining regions. The catalytically important C residue is the first from J4/2 (C_57_). **D**) Schematic representation of the CPEB3 ribozyme dimer as interpreted from the crystal structures. The upper inset indicates the actual secondary structure taking place in the human J1/2 region.

In the following years, structures of uncleaved (Chen et al. 2010), mutant-inhibited (Kapral et al. 2014), and ion-complexed forms (Ke et al. 2004; Ke et al. 2007) were published, confirming that both pre- and post-cleavage states of the genomic HDV ribozyme, adopt nearly identical folds, including the nested double pseudoknot. The catalytic core of the HDV ribozyme is flanked by two highly conserved G•U wobble base pairs (bp) playing different structural functions. The first one corresponds to the ribozyme’s cleavage site at the first bp of P1. This *cis* G1•U37 wobble has a local stabilizing effect on the hydrogen bonds between G1 and the critical catalytic residue C75 and has been shown to bind Mg^2+^ (Cerrone-Szakal et al. 2008a; Cerrone-Szakal et al. 2008b), as does the second wobble bp located in P3, the *trans* G25oU20 (Chen et al. 2010). This bp conformation slightly adapts to its dynamic environment during self-cleavage and has a stabilizing action on P1.1 (Sripathi et al. 2015).

In 2006, using a human transcriptomic library designed for searching ribozymes, the Szostak group discovered a new self-cleaving ribozyme located in the second intron in the pre-mRNA encoding the cytoplasmic polyadenylation element binding 3 protein (CPEB3) (Salehi-Ashtiani et al. 2006) (**Figure 1B**). Interestingly, HDV-like ribozymes are found in a wide variety of organisms (reviewed in (Webb and Luptak 2011)). Based on biochemical and secondary structure characterization, this ribozyme was classified as the first representative of the HDV-like ribozyme family (Webb and Luptak 2011) (**Figure 1C**). The *CPEB3* gene encodes the prion-like CPEB3 protein, a protein involved in synaptic plasticity and long-term memory (Fioriti et al. 2015; Stephan et al. 2015; Ford et al. 2019; Ford et al. 2023). The ribozyme sequence is highly conserved among mammals, and the self-cleavage activity has persisted over > 170 million years of evolution (Bendixsen et al. 2021). However, despite its apparent similarity to the HDV ribozyme, very little is known about the ribozyme mechanism and function. It has been suggested that coupling between splicing and ribozyme self-cleavage may regulate the CPEB3 mRNA/protein levels and thus have an impact on synaptic plasticity (Chen et al. 2021).

Here, we focus on the human and chimpanzee homologs of the CPEB3 ribozyme. The sequences of these ribozymes differ only at position 30, which seems sufficient to make the chimpanzee ribozyme cleave faster than the human one (**Figure 1C**) (Chadalavada et al. 2010; Bendixsen et al. 2021). Comparative sequence analysis suggests that P1 from the human CPEB3 ribozyme is closed by a wobble C•A pair (also found in sloth, armadillo, and dolphin), while other mammals, including the chimpanzee, present a *cis* Watson-Crick C-G bp (Webb and Luptak 2011).

To investigate the three-dimensional structure of the CPEB3 family members, we crystallized the cleavage products of the human and chimpanzee CPEB3 ribozymes and report here their crystal structures. Both structures confirm the general topology observed in the HDV ribozyme with two coaxial helical stacks. However, we observe an unforeseen conformation where the P1.1 strands are not base paired, in opposition with NMR data (Skilandat et al. 2016). This opening allows for forming a four bp stem between two L3 loops of two symmetry-related CPEB3 ribozymes (**Figure 1D**). Strikingly, the base paired 7 nucleotides (nt) L3 loops fold in a conformation similar to the one from the anticodon loop (AC) of tRNAs when interacting with the mRNA codon. The dimerization is promoted by the four nucleotides palindrome downstream from the U-turn, free to base pair with a neighbouring RNA molecule. Interestingly, RNA dimerization can either be considered as a crystallization artefact as in the case of tetraloop-containing oligonucleotides (Holbrook et al. 1991; Baeyens et al. 1995; Baeyens et al. 1996; Lietzke et al. 1996) or in the case of the HIV dimerization insertion sequence (Ennifar et al. 1999; Ennifar et al. 2001). Nevertheless, ribozyme dimerization can also be biologically relevant, as in the case of the Varkud satellite ribozyme (Suslov et al. 2015), the hammerhead ribozymes from Penelope-like transposable elements (Lunse et al. 2017), as well as the hatchet ribozyme (Zheng et al. 2019).

A precise structural analysis of the genomic, antigenomic HDV, and CPEB3 ribozymes indicates that the antigenomic HDV ribozyme presents conserved features from both the CPEB3 ribozyme and the genomic HDV ribozyme. This work supports the hypothesis made by Szostak that the HDV ribozymes may have evolved from the CPEB3 ribozyme (Salehi-Ashtiani et al. 2006).

## RESULTS

### Design of the CPEB3 ribozyme constructs

Based on pioneering work from Ferré-D’Amaré and colleagues (Ferré-D’Amaré et al. 1998a), CPEB3 ribozymes harboring a nine nt leader sequence 5’-GGGAUAUCA-3’ were produced by *in vitro* transcription (IVT). The cleavage products were separated from those trapped in non-active conformations by denaturing PAGE. To promote crystal packing interactions, the non-essential P4 region was replaced by a U1A protein RNA binding domain (RBD) (Ferré-D’Amaré et al. 1998b; Rupert and Ferre-D’Amare 2001; Cochrane et al. 2007; Ferre-D’Amare 2010; Kulshina et al. 2010; Reiter et al. 2010; Costa et al. 2016). Accordingly, both human and chimpanzee CPEB3 ribozymes were produced with the U1A-RBD substitution in the catalytically neutral P4 helix (**Figure 1C**). The effectiveness of the cleavage reaction in the course of the IVT reaction indicates that the insertion of the U1A binding site has no inhibitory effect on the ribozyme (**Figure S1**). Prior to crystallization, the formation of the RNA/U1A protein complex was assessed by electrophoretic mobility shift assay (EMSA) (**Figure S2**). The present crystal structures unravel for the first time the human and the chimpanzee CPEB3 ribozymes 5’-hydroxyl product state, captured as dimers.

### A L3 loop sequence compatible with the formation of an anticodon-like conformation favors CPEB3 ribozyme dimerization over folding of a catalytically competent conformation

At first glance, the 2D structures of the HDV and CPEB3 ribozymes look very similar. However, the sequence determinants and ribozyme length differ (**Figure 1A and C**). The minimum HDV ribozyme (PDB ID: 4pr6) is about 85 nt (Perrotta and Been 1991) versus 67 nt for the CPEB3 RNA with the wild-type P4. Interestingly, the most important differences are located around the catalytic core: (i) the CPEB3 ribozyme putative P1.1 mini-helix consists of one canonical G-C bp and one U•U pair instead of two canonical G-C bp in the HDV ribozyme; (ii) the 3 nt joining region J1.1/4 is absent in the CPEB3 ribozymes; and (iii) the 5’-A_23_CGU_26_-3’ stretch of L3 presents 4 nt versus 5 nt (including only one purine) in the genomic HDV ribozyme; (iv) P4 in the CPEB3 ribozyme is shorter than in the genomic HDV ribozyme. Although NMR studies mainly point to a ribozyme monomer in solution (Skilandat et al. 2016), the dimerization of the CPEB3 ribozymes observed in this study seem to result from these sequence and secondary structure changes, which affect ribozyme folding, 3D structure, and cleavage rate (**Figure 2A**).

**Figure 2.**
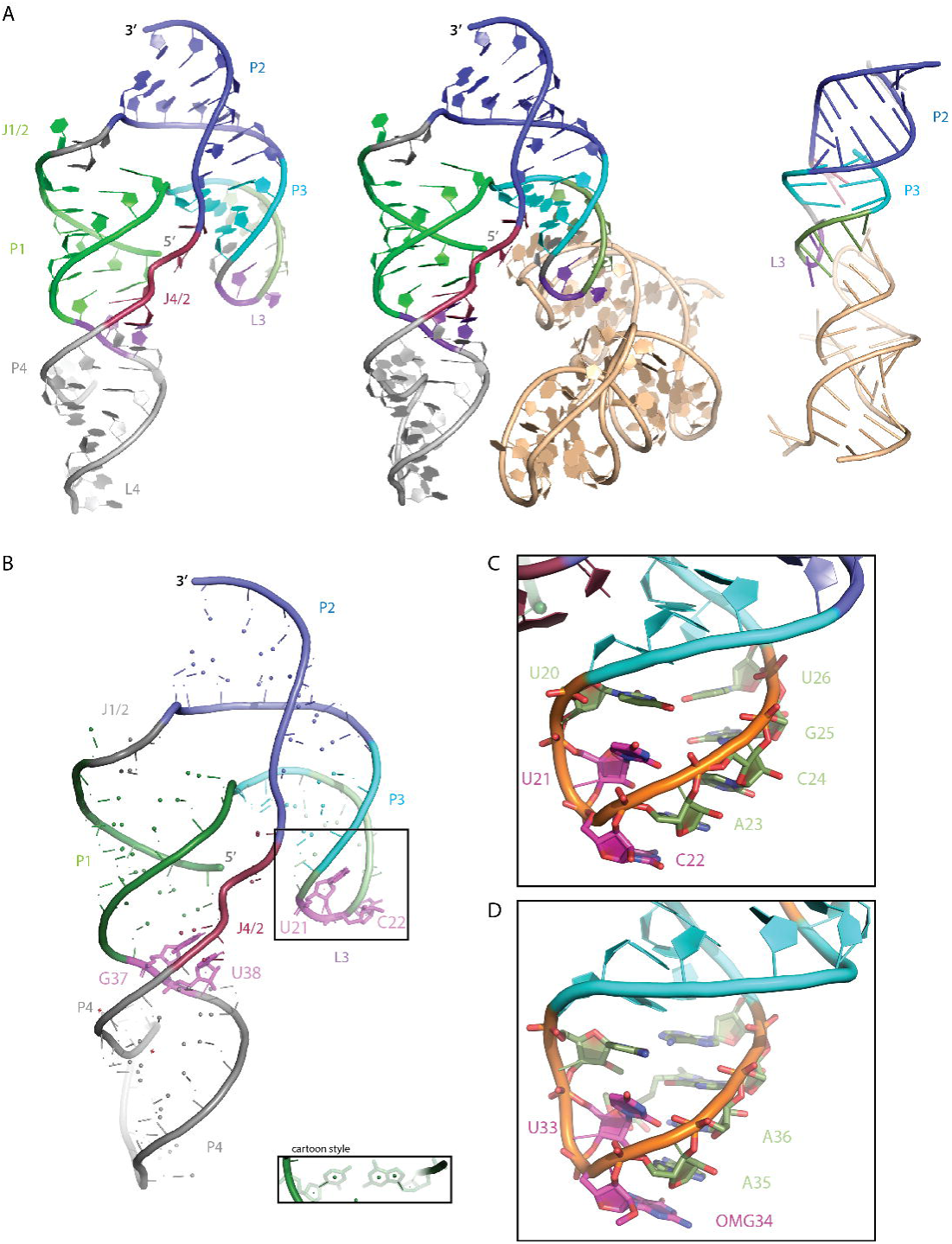
Crystal structures of the human (left) and the chimpanzee (right) CPEB3-U1A-RBD ribozymes. (A) The human CPEB3-U1A ribozyme (left panel) is a crystallographic dimer (central panel) while the chimpanzee CPEB3-U1A is pseudo-symmetric. The dimerization site (right panel) comprises nucleotides A23, C24, G25, and U26 from the L3 loop. The U1A protein binding loop L4 is omitted for clarity. (B) The residues predicted to form the mini-helix P1.1 (purple) are highlighted in sticks on the cartoon 3D backbone spline. The mode of representation is illustrated in the inset entitled “Cartoon style”. (C) shows a close-up of L3 highlighting residues supposed to form the nested double pseudoknot (U21, C22, purple) and the five residues involved in dimerization (C22, A23, C24, G25 and U26, green). The nucleobases of U20 and U26 are coplanar but could only form water-bridging hydrogen bonds deduced from the distance between the two proximal carbonyl groups. A23 to U26 adopt a helical conformation. U21 adopts a classical U-turn conformation and its N3 contacts the O2P oxygen of C24. (D) The anticodon loop of tRNAPhe (PDB ID: 1ehz (Shi and Moore 2000)) is depicted for a direct visual comparison, which indicates that each nucleotide conformation is similar to the one observed in L3 from the CPEB3 ribozyme.

In the observed conformations of the CPEB3 ribozymes, the 7 nt L3 loop residues U_21_ and C_22_ are splayed apart by the U-turn, hence prevented to pair with their respective targets U_38_ and G_37_ to form the P1.1 mini-helix (**Figure 2**). This feature is accompanied by the formation of an intermolecular 4 bp helix taking place between the L3 palindromic stretches (5’-A_23_CGU_26_-3’) of two (pseudo)symmetry-related ribozymes. The positions of these residues within the loop partially correspond to those forming the codon-anticodon interaction during ribosome translation. In this unforeseen ribozyme dimer, a competent catalytic core cannot be formed. The dimerization could arise from the competition between a weak intramolecular two bp P1.1 mini-helix (formed by one G-C bp and one weaker U•U pair) and a stronger intermolecular four bp helix formed by nucleotides 5’-A_23_CGU_26_-3’ from L3. The dimerization interaction propagates through all L3 residues adopting a helical conformation, including C_22_, which forms a loose triple interaction with U_20_ and U_26_ from a symmetry-related molecule. Some contacts are also observed between two U1A proteins, which may contribute to the further stabilization of the dimer. However, RNA crystal structures obtained with the help of the U1A RBD usually present U1A-mediated crystal packing contacts that do not interfere with the RNA global fold (Ferre-D’Amare 2010).

Consequently, the G_37_ and U_38_ residues from J1/4 supposed to form P1.1 are sequestered by those from J4/2, resulting in extending P4. These two non-canonical additional base pairs mediate stacking continuity with P1 (**Figure 2B**), and also prevent the assembly of the catalytic site. Despite CPEB3 ribozyme dimerization, the type I A-minor interaction (Doherty et al. 2001; Nissen et al. 2001) involving A_62_ and the P3 penultimate bp (C_18_-G_28_) is observed (corresponding to A_77_sC_18_wcG_29_ in HDV (PDB ID: 1drz)) whereas the type II that would be expected from the catalytic site assembly observed in the HDV ribozyme structure (A_76_sC_19_wcG_28_) is dissociated (“s” stems from the sugar edge and “wc” from Watson-Crick (Leontis and Westhof 2001)). The HDV ribozyme catalytic core is flanked by the *cis* G1•U37 and the *trans* G25oU20 wobble pairs (HDV numbering, **Figure 1A**), which are crucial for ribozyme activity since their mutations significantly reduce cleavage rate. In the crystal structures of the CPEB3 ribozyme the G25oU20 is disrupted by ribozyme dimerization.

Another notable difference between all HDV ribozymes is the presence of the three nt J1.1/4 stretch only in the genomic ribozyme (**Figure 1A**). There, P4 starts by a A_43_wcG_74_ bp, and J1.1/4 adopts a specific turn, which resembles an AA-platform (the sugar and Hoogsteen edges of G_40_ and A_42_, respectively, form hydrogen bonds) (Cate et al. 1996) with a bulging C inserted. Moreover, the WC edge from G_40_ interacts with the Hoogsteen edge of G_74_. The area mediated by this triple interaction provides a stacking surface for both P1.1 on one side and P4 on the other. The stacking continuity between the P1, P1.1 and P4 elements thus mainly results from the presence of J1.1/4. The absence of J1.1/4 raises the question of how P1.1 can be stabilized in the antigenomic HDV, and especially in CPEB3 ribozymes considering the intrinsic weakness of the P1.1 U-U bp. In the present crystal structures of the CPEB3 ribozymes, the stacking continuity between P1 and P4 relies on residues G_37_ and U_38_, which are embedded in P1.1 in a catalytically competent conformation (Skilandat et al. 2016; Roberts et al. 2023). How these residues could be untrapped to reach a catalytically competent conformation is not yet understood.

### A single nucleotide change between the human and chimpanzee CPEB3 ribozymes structures supports their different catalytic properties

A unique sequence difference between the human (A_30_) and the chimpanzee (G_30_) homologues induces a local structural adaptation (**Figure 3 and 4**). The putative C•A pair proposed by comparative sequence analysis to close P1 in the human CPEB3 ribozyme does not form. Instead, C_7_ is pushed away towards the solvent and its presumed partner A_30_ is in close distance to A_8_(O2’) (**Figure 4**) - setting up this conformation led to a significant decrease of R_free_ from 0.2870 to 0.2602. The proximity of A_30_ to O2’ and possible stacking on G_31_ obviously helps stabilizing J1/2. As C_7_ is pushed away, C_6_, A_8_ and C_9_ stack one on each other, positioning A_10_ to form a standard WC bp with U_69_ (**Figure 4b**). Per contra, the chimpanzee CPEB3 ribozyme conserves the C7-G30 canonical bp, smoothly curving the backbone, placing all nucleobases towards P2 (**Figure 4a**). Consequently, the shorter J1/2 of the human version of the ribozyme leads to a shorter distance between the proximal ends of P1 and P2 than in the chimpanzee ribozyme (d_chimp_(O3’ (C7)-P(A10)=12.2 Å; d_hum_(O3’ (A8)-P(A10)=7.4 Å). The regularity of the human P1 is broken by the non-canonical A-A pair and the bulge, potentially affecting its stability. Otherwise, the overall ribozyme conformations are very similar, conserving the two parallel coaxial helical stacks described earlier, and the same dimer orientation.

**Figure 3.**
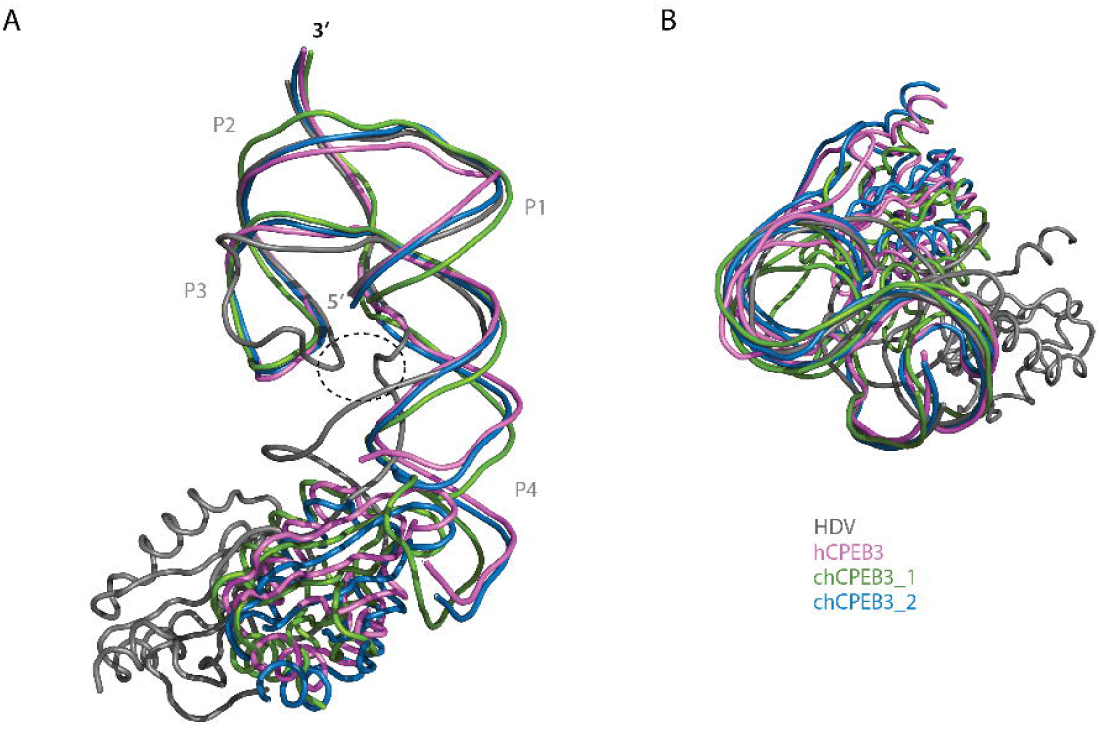
Structural similarity between the CPEB3 and the HDV ribozymes. Alignment of the genomic HDV (PDB ID: 4PR6), of the human CPEB3-U1A ribozyme (PDB ID: 7qr4) and of the chimpanzee CPEB3-U1A (PDB ID: 7qr3) ribozymes. Both conformations of the chimpanzee CPEB3-U1A ribozymes are represented. The HDV ribozyme is shown in gray, the human CPEB3-U1Ar in violet and both chimpanzee CPEB3-U1Ar conformations in green and blue. The dashed ellipsoid delineates the shorter distance taking place in the genomic HDV ribozyme structure due to the formation of P1.1.

**Figure 4.**
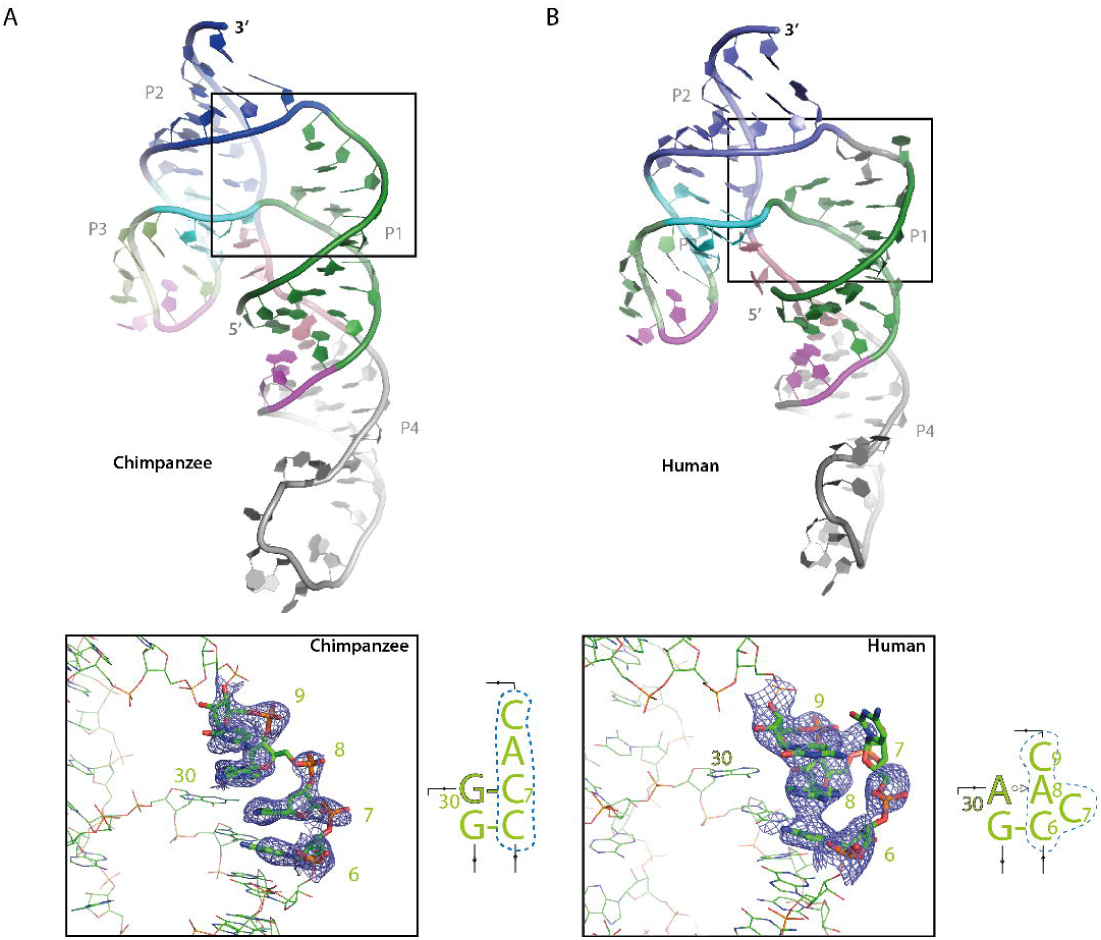
Structural differences between human and chimpanzee CPEB3 ribozymes around J1/2 at the P1/ P2 strand cross-over. Both chimpanzee (A) and human (B) CPEB3 ribozymes present the same J1/2 sequences, although they are structured differently due to a sequence difference at position 30 (outlined nucleobase). A30 favours a non-canonical A30w_o_sA base pair instead of a G30-C base pair. Consequently, residues 6 to 9 form a stacked column in the chimpanzee ribozyme, whereas in the human ribozyme, only 3 residues can stack with C7 bulging out. The distance between the proximal extremities of P1 and P2 is thus reduced to less than 10 Å in the human ribozyme. The weighted 2Fobs-Fcalc electron density maps are drawn contoured at 1.0 sigma. The corresponding region is highlighted by a blue dashed frame on the secondary structure diagrams corresponding to each situation.

## DISCUSSION

The genomic HDV ribozyme has been the subject of intensive biochemical studies over years. The first crystal structure of the ribozyme cleavage product was published over two decades ago (Ferré-D’Amaré et al. 1998a) followed by the pre-cleavage form in the presence of various ions (Ke et al. 2004; Ke et al. 2007) and an inhibited trans-acting ribozyme form (Chen et al. 2010). Here, we present the crystal structures of the self-cleavage products of both the human and the chimpanzee HDV-like CPEB3 ribozymes. As in the case of the historic HDV crystal structure, the P4 region of both ribozymes were replaced by the U1A RNA binding domain, a well-known effective crystallization module (Ferre-D’Amare 2010). In the absence of structural data on the CPEB3 ribozymes family, the initial secondary structure integrated the tertiary contacts observed in the HDV ribozyme, including the stacking between P1 and P4 on one side and P2 and P3 on the other side, with the formation of the P1.1 two bp mini-helix resulting from the interaction between the L3 loop and the residues tethering P1 to P4 (J1/4). Yet, even though the stacking partners remain, the conformers significantly depart from the HDV archetype, notably due to the opening of P1.1, which promotes dimerization through the interactions of the L3 loops of two ribozymes. The L3 loops unexpectedly acquire a tRNA anticodon loop-like shape. These unforeseen conformations for HDV-like ribozymes, indeed expands the structural repertoire of the HDV ribozyme family.

It has been suggested that the chimpanzee ribozyme is more prone to adopt its native state due to a more stable P1 helix than its human counterpart (Chadalavada et al. 2010). Indeed, the regular P1 stem may better stabilize the chimpanzee ribozyme than the human ribozyme, where a bulge (C_7_) followed by a WC-sugar edge A-A pair breaks P1 regularity (**Figure 1D and 4B**). Consequently, the difference between the two CPEB3 ribozyme homologues may influence ribozyme compaction and further their catalytic activities.

Although the CPEB3 ribozyme is highly conserved in mammals and its localization within an intronic sequence has remained unchanged for 170 million years (Bendixsen et al. 2021), its precise mechanism of action and function remain elusive. Both *Aplysia* and *Drosophila* CPEB3 homologues (CPEB and ORB2, respectively) have been shown to play a crucial role in neuronal plasticity and their function strongly correlates with their ability to form prion-like oligomers (Hervas et al. 2021). In humans, the only CPEB protein able to aggregate is CPEB3. Work from Kandel’s laboratory has shown that the human CPEB3 protein is necessary for synaptic plasticity by stimulating cytoplasmic polyadenylation-induced translation of plasticity related proteins (PRPs) like AMPA (α-amino-3-hydroxy-5-methyl-4-izoazolepropionic acid) receptor subunits GluA1 and GluA2 (Fioriti et al. 2015; Ford et al. 2019). CPEB3 exists in both soluble and aggregated states depending on SUMOylation, having a regulatory effect on the translation of several mRNAs (Drisaldi et al. 2015; Ford et al. 2023). However, the link between the CPEB3 ribozyme activity, the protein expression level, and the hippocampal-dependent memory remain to be elucidated. A recent study from the Luptak laboratory has shown that antisense oligonucleotides (ASO) competing with P1 formation lead *in vitro* and in vivo to increased levels of both the CPEB3 protein and its mRNA, presumably by perturbing CPEB3 ribozyme folding and catalysis (Chen et al. 2021). The authors conclude that the ribozyme activity influences the expression of CPEB3, which confirmed the original hypothesis from Salehi-Ashtiani et al. (Salehi-Ashtiani et al. 2006) that the low background activity is linked to splicing completion.

The present crystal structures display inhibitory conformations of the ribozymes’ cleavage products. It is thus possible to directly compare the structural roles of the individual nucleotides according to their identity and position in the CPEB3 and HDV ribozymes. Interestingly, in the present conformers, the folding of L3 in an anticodon-like loop conformation with the concomitant sequestration of the J1/4 residues by those from J4/2, including the catalytic C_57_, would be sufficient to repress catalytic activity, by preventing the formation of the P1.1 mini-helix. Nevertheless, the anticodon-like loop is poised to dimerize through the formation of a 4 bp double-stranded helix (**Figure 2**). Even though we cannot infer from the structures whether dimerization is biologically relevant, we can suggest that the potential to do so would significantly perturb the equilibrium between the open and the catalytically active conformation. Two sequence elements seem to drive dimerization, the palindrome (5’-ACGU-3’) corresponding to the tRNA anticodon nucleotides 35 to 38 (tRNA numbering). C_22_ upstream from the palindrome (corresponding to residue 34 in tRNA numbering) contributes a loose triple interaction with the first and the ultimate residues of L3, U_20_ and U_26_ of the interacting RNA. The second important element is U_21_ corresponding to the almost fully conserved U_33_ (tRNA numbering), which mediates the U-turn typical of tRNAs anticodon loops (Pak et al. 2017). This residue is also observed to form the weak UoU pair involved in the P1.1 pseudoknot by high-throughput pairwise mutational studies (Roberts et al. 2023) or NMR (Skilandat et al. 2016). The conserved 4 nt palindrome together with the dual role of U_21_ may favor the inactive conformation of the CPEB3 ribozyme through dimerization.

It has been proposed by the Szostak group that the HDV ribozymes have originated relatively recently from the CPEB3 ribozyme encoded by the human transcriptome (Salehi-Ashtiani et al. 2006). In a distinct genetic context, the release of the constraints due to coupling to splicing may thus have allowed the sequence of the CPEB3 ribozyme to drift up to reach faster cleavage properties gathered in what are known today as HDV ribozymes. Such a phenomenon has been demonstrated in the case of the lariat-capping ribozyme, a ribozyme structurally related to group I introns (Meyer et al. 2014). To achieve this, the HDV ribozymes may have gradually lost the propensity to fold L3 as an anticodon-like loop prone to dimerize and compete with building a cleavage-competent conformation without being kinetically trapped. These drifting events included (i) mutation of the U-turn resulting in strengthening the P1.1 pseudoknot required for building up the catalytic pocket, (ii) mutations breaking down the palindrome. Beyond these changes, the genomic HDV ribozyme intermediates may have acquired (iii) an additional nucleotide in L3 and (iv) a J1.1/4 stretch that should also improve its catalytic properties. In between these two extreme points stands the antigenomic HDV ribozyme, which has not yet gained the J1.1/4 junction, yet still harboring a non self-complementary Y-rich 7 nt L3 loop (**Figure 5**). In spite of cleavage constants in the same range (∼1 min^-1^) for the minimal forms of the genomic and antigenomic HDV ribozymes, their different genetic context may have influenced their sequence specificities (Brown et al. 2008). The present structures captured a state in which P1.1 is actually molten. This feature is observed by X-ray crystallography herein for the first time. Melting of P1.1 could result from a less stable base pair scheme conferring poor catalytic properties to the CPEB3 ribozymes. This observation has to be put in the context of the dual roles of the P1.1 residues embedded in L3, which competitively adopts an anticodon-like loop conformation poised to dimerize by means of a palindromic four nt sequence. However, the present results do not rule out whether the dimers observed in the human and chimpanzee CPEB3 ribozymes have any biological relevance.

**Figure 5.**
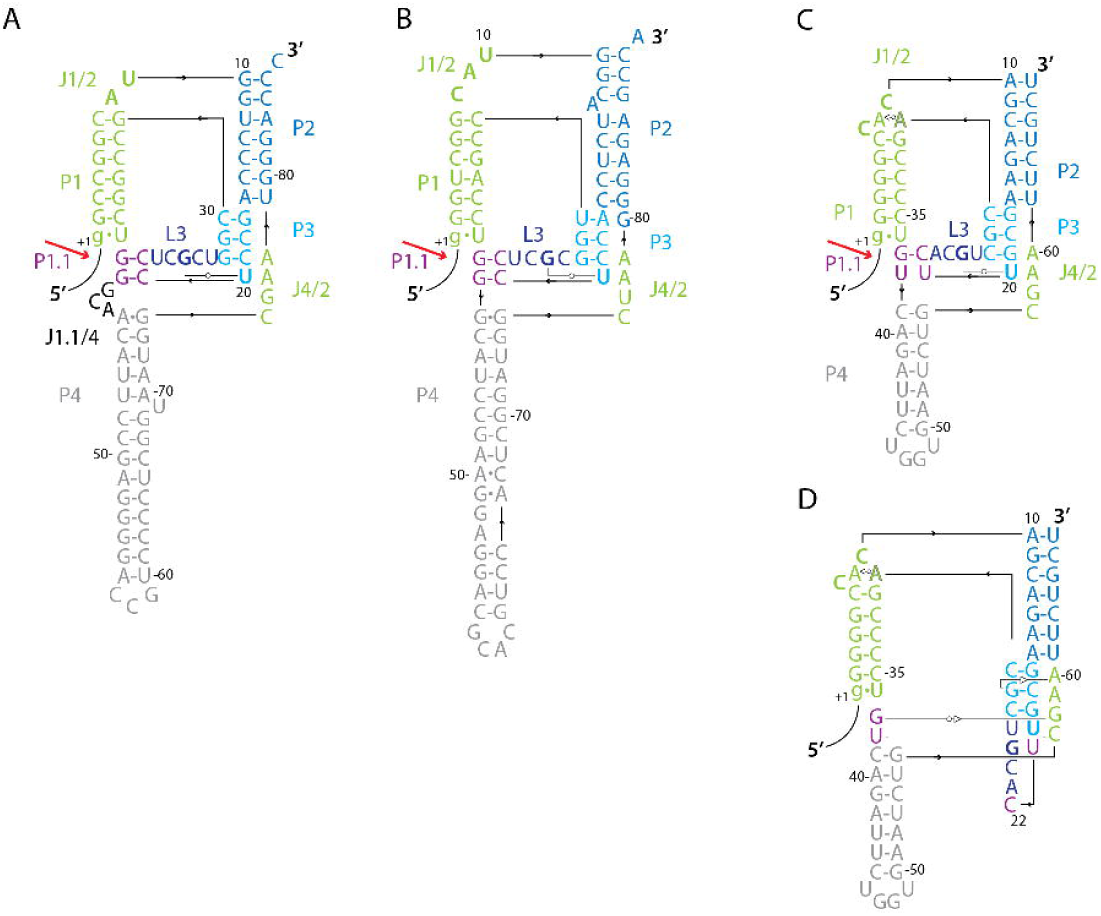
Secondary structures of the HDV and CPEB3 ribozymes. The differences between these ribozymes mostly reside in L3 (deep blue), P1.1 (purple), and J1.1/4 (black). (A) The genomic HDV ribozyme presents an 8-membered loop not expected to adopt an anticodon conformation. P1.1 is composed of two GC pairs with a C_21_ residue not supporting the appearance of a U-turn. The J1.1/4 junction supports the formation of P1.1. (B) The antigenomic HDV ribozyme has no palindrome and is Y-rich. L3 still has 7 residues. Yet the presence of C_21_ is not expected to support the U-turn conformation of L3. (C and D) The CPB3 ribozyme presents a seven membered L3 loop containing a four nt palindrome 5’-ACGU-3’. U21 supports the formation of a U-turn inducing competition between the formation of an antidcodon-like conformation for L3 (D) and the closure of P1.1 to drive the ribozyme towards catalysis (C).

Nevertheless, other studies by the Bevilacqua laboratory have pointed out the multimerization of the human CPEB3 ribozyme. Strulson et al. (Strulson et al. 2013) have reported the detection of human CPEB3 ribozyme aggregates by PAGE under native conditions in the presence of high molecular weight crowding agents and 10 mM [Mg^2+^] - indeed, looking carefully at figure S7 of the study from Strulson (Strulson et al. 2013) suggests that the band interpreted as aggregated CPEB3 ribozymes could correspond to dimers. Chadalavada et al. (Chadalavada et al. 2010) have also observed the precursor of the CPEB3 by PAGE native conditions and have noted two distinct species they called R1 and R2. These species are separated one from each other on native gels so that they could correspond to dimer and monomer in the light of the present crystal structures (See figure 2 of (Chadalavada et al. 2010)).

The question we will need to address in the future is whether the present dimeric structures result from a crystallization artefact. A number of studies point to ribozyme dimers as functional entities. In 2017, numerous (±150 000) small hammerhead ribozymes have been discovered characterized by a short (1 bp) stem III and relatively poor cleavage rate (Lunse et al. 2017). These ribozymes become active upon forming dimers presenting a recapitulated stem III. The VS ribozyme has also been shown to be active as a dimer in its natural context (Suslov et al. 2015). The Hatchet ribozyme has also been crystallized as a dimer of cleavage products (Zheng et al. 2019). In this case, the 3’ tail of one ribozyme entangles the core of a pseudo-symmetric ribozyme and vice-versa. Strikingly, the entangling sequence contains the same palindrome 5’-ACGU-3’ as the CPEB3 ribozymes (Bou-Nader and Zhang 2020). All these data point towards the necessity to investigate the biological significance of the inactive CPEB3 ribozymes conformations prone to dimerize in order to better understand the relationship between the CPEB3 ribozyme activity and the expression of the CPEB3 protein.

## MATERIAL AND METHODS

### RNA preparation

The DNA templates were designed to encode the human or chimpanzee CPEB3 ribozyme sequence with a U1A-RBD within P4 and a 9 nt 5’ leader sequence to allow self-scission of the ribozyme during transcription. *In vitro* transcription reactions (Gallo et al. 2005) were performed using 0.85 µM of the DNA template strands (Microsynth), 5 mM of each NTP (GE Healthcare), 0.01% Triton X-100, 35 mM MgCl_2_, 200 mM Tris-HCl pH 7.5, 200 mM DTT, 10 mM Spermidine, and T7 polymerase and were incubated for 4 hours at 38°C. The RNA was purified by 12% denaturing PAGE, visualized by UV shadowing, and recovered by electroelution, desalted using Vivaspin^®^ concentrators (5000 Da MWCO (Viva products)). The RNA was resuspended in 100 mM KCl, 10 mM HEPES-KOH pH 7.5, 10 µM EDTA buffer at 0.4 mM.

### Double-Mutant U1A protein

The double-mutant U1A protein plasmid was a kind gift from Adrian Ferré-d’Amaré. The protein was produced as described by Ferré-d’Amaré (Ferré-D’Amaré et al. 1998b), omitting the last hydroxyapatite (CHT-I, BioRad) chromatography step. SDS-PAGE and mass spectrometry confirmed the final product.

### Complex formation

The purified RNA was refolded by 1 minute incubation at 90°C followed by 4 minutes on ice. The refolded RNA was annealed to the Double-Mutant U1A protein 1:1 molar equivalent ratio to yield a complex concentration of 0.2 mM (in 50 mM KCl, 6.7 mM HEPES-KOH pH 7.5, 5 nM EDTA, 5 mM MgCl_2_, and 1 mM spermine).

### Crystallization

For crystallization by the vapor-diffusion technique, 400 nL sitting drops of complex solution were mixed with 200 nL of reservoir solution. The initial crystals of both complexes were obtained at the Protein Crystallization Center (PCC, UZH) using a Gryphon LCP robot (ART - Art Robbins Instruments) and the Index HT sparse matrix screen (Hampton Research) for the human CPEB3 complex and PEG/Ion screen (Hampton Research) for the chimpanzee complex. The human CPEB3 complex was crystallized in a solution containing 20-24% PEG 3350, 0.2 M sodium citrate tribasic dihydrate and 30-200 mM HEPES-KOH pH 7.5. For the chimpanzee CPEB3 complex, the reservoir solution contained 19-24% PEG 3350, 0.2 M sodium formate. The crystallization assays were incubated at 4°C. The human and chimpanzee CPEB3 complex crystals appeared within 5 days, and grew over 25 days, and reached maximum dimensions of 250 x 50^2^ µm^3^. For cryogenic conditions, the crystals of the human CPEB3 complex were transferred to cryoprotecting solution containing 25% PEG3350, 10 mM HEPES-KOH pH 7.5, 100 mM KCl, 10 mM MgCl_2_, 2 mM spermine, 0.2 M sodium citrate tribasic dihydrate, and 5% or 10 % (v/v) glycerol. The crystals of the chimpanzee CPEB3 complex were transferred to a solution containing 22% PEG3350, 10 mM HEPES-KOH, pH 7.5, 100 mM KCl, 10 mM MgCl_2_, 2 mM spermine, 0.2 M sodium formate, 10% or 18% (v/v) glycerol. Crystals were mounted into CryoCap loops (Molecular Dimensions) and flash-cooled in liquid nitrogen.

### Data collection, structure determination and refinement

Native datasets were collected at the beamline PXIII (X06DA) at the Paul Scherrer Institute (PSI), using a Pilatus 2M-F detector and the multi-axis PRIGo goniometer at 1.00004 Å wavelength with an exposure of 0.2 s per 0.2° for 360° data set.

Data were indexed in space group C222_1_ and processed using the autoPROC pipeline. Diffraction frames were integrated by XDS and scaled by Aimless (Evans and Murshudov 2013). Anisotropy was corrected by Staraniso (Bricogne et al. 2011). The human complex structure was solved by the molecular replacement method in Phaser (McCoy et al. 2007), using 4pr6 as a search model. Missing nucleotides were built by hand. The chimpanzee complex structure was solved using the human complex as a template and anomalous data were collected with X-ray energy at 5.975 keV to localize phosphorus and sulfur atoms. Iterative model building and refinement was performed using PHENIX (Adams et al. 2010), COOT (Emsley et al. 2010), and Buster 2.10.3 (Blanc et al. 2004). More than 370 crystals were screened to solve the human CPEB3 ribozyme structure, owing to the temperature sensitivity of the crystals and a high solvent content of 74%. The highest resolution was 2.83 Å. Only 20 crystals were screened to determine the chimpanzee CPEB3 ribozyme structure. They had a solvent content of 62% and diffracted to a resolution of 2.18 Å. The human CPBE3-U1A ribozyme is a crystallographic dimer while the chimpanzee ribozyme is pseudo-symmetric. The refined models were validated with MolProbity (Davis et al. 2004). Data collection and refinement statistics are shown in **Supplementary Table 1**. All figures were prepared using PyMOL (Schrodinger 2010).

### Pair Fit Alignment

Root Mean Square Deviation (RMSD) determination was performed by the Pair Fit Alignment tool in PyMol (Schrodinger 2010). The phosphorus atoms of the nucleotide stretches 101 to 122 and 163 to 172 of the genomic HDV structure (PDB ID: 4pr6) were aligned with the nucleotide stretches 1 to 22 and 59-68 for both the human and chimpanzee CPEB3 ribozymes. These residue ranges were chosen to identify structural differences between the HDV and CPEB3 ribozymes without focusing on the P4 and U1A-RBD domains. Alignment of the three CPEB3 ribozyme structures was based on their full RNA sequence (nucleotides 1 to 69).

### Sequences

provided in DNA Short template (forward strand):

5’-GAAATTAATACGACTCACTATAGG-3’

Human-CPEB3-U1A ribozyme (reverse strand):

5’-AGCAGAATTCGCAGCCGGAGTGCAATGGCTGACAGGGGCTGCGACGTGAACGCTTCTGCTGTGGCC CCC<u>TGTTATCCC</u>**TATAGTGAGTCGTATTAATTTC**-3’

Chimpanzee CPEB3-U1A ribozyme (reverse strand):

5’-AGCAGAATTCGCAGCCGGAGTGCAATGGCTGACAGGGGCCGCGACGTGAACGCTTCTGCTGTGGCCCCCTGTTATCCCTATAGTGAGTCGTATTAATTTC-3’

## ACCESSION NUMBERS

Crystal structure models and structure factors were deposited at the wwPDB. PDB IDs are 7qr3 for the chimpanzee and 7qr4 for the human CPEB3 ribozymes.

## ACKNOWLEDGMENT

This work was supported by the fund of the Forschungskredit (FK-18-097) of the University of Zürich to A.I.P.M and the Swiss National Science Foundation (projects 200020_165868 and 200020_192153) to R.K.O.S. B.M. was supported by the Interdisciplinary Thematic Institute IMCBio, as part of the ITI 2021-2028 program at the University of Strasbourg, CNRS and Inserm, by IdEx Unistra (ANR-10-IDEX-0002), and EUR (IMCBio ANR-17-EUR-0023), under the framework of the French Investments Program for the Future. We also thank Dr. Adrian Ferré-D’Amaré for the kind gift of the double mutant-U1A protein plasmid and Prof. Eva Freisinger for her help handling the crystals.

