## Supplementary figures and tables for "Anticodon-like loop-mediated dimerization in the crystal structures of HDV-like CPEB3 ribozymes"

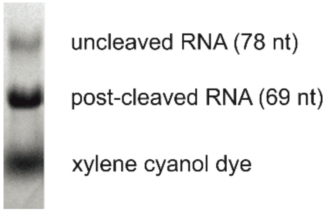

**Figure S1. Cotranscriptional self-cleavage.** Representative 12% denaturing PAGE showing 5'-end self-cleavage of human CPEB3-U1A ribozyme during transcription. The *in vitro* transcription/cleavage reaction was performed at 37°C for 2 hours.

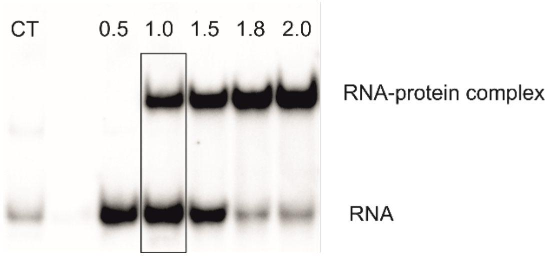

**Figure S2. Formation of an RNA-protein complex.** Electrophoretic mobility shift assay (EMSA) of [α-$^{32}$P]-CTP labeled human CPEB3-U1A ribozyme titrated with DM-U1A protein. Samples contained $2.87 \cdot 10^{-7}$ M RNA and 0 (CT), 0.5, 1, 1.5, 1.8, and 2.0 eq DM-U1A protein, respectively, in 100 mM KCl, 5 mM MgCl2, 1 mM spermine and 10 mM HEPES-KOH at pH 7.5. The 1:1 ratio used for crystallization is marked with a black rectangle.

**Table S1. Data collection and model refinement statistics.**

| Data collection | Human CPEB3 ribozyme | Chimpanzee CPEB3 ribozyme |
|---|---|---|
| Space group | $C222_1$ | $C222_1$ |
| Unit cell parameters *(a; b; c (Å))* | 79.451; 131.879; 90.456 | 120.254; 135.976; 83.254 |
| Resolution range *(Å)* | 68.055-2.825 (3.124-2.825)[a] | 67.958-2.184 (2.355-2.184)[a] |
| Total no. of reflections | 87407 (4303)[a] | 374221 (19001)[a] |
| Unique reflections | 6828 (342)[a] | 28178 (1410)[a] |
| Completeness *(%) (ellipsoidal)* | 92.6 (73.7)[a] | 93.7 (57.0)[a] |
| $R_{merge}$ *(all $I^+$ and $I^-$)* | 0.081 (1.998)[a] | 0.064 (1.904)[a] |
| $R_{meas}$ *(all $I^+$ and $I^-$)* | 0.085 (2.083)[a] | 0.067 (1.985)[a] |
| Multiplicity | 12.8 | 13.3 |
| *MeanI/σI* | 16.7 | 24.9 |
| $CC_{1/2}$ | 0.998 | 1.000 |
| **Models** | | |
| PDB number | 7qr4 | 7qr3 |
| $R/R_{free}$ | 0.220/0.268 | 0.241/0.277 |
| No. Reflections used to $R_{free}$ | 348 | 1431 |
| Wilson *B* factor | 93.31 | 57.16 |
| Number of atoms<br>RNA<br>Protein | <br>1461 (chain B)<br>728 (chain A) | <br>1462/1462 (chains C/D)<br>738/742 (chains A/B) |
| Rmsd from ideal values<br>Bond lengths (Å)<br>Bond angles (°)<br>Dihedral angles (°) | <br>0.003<br>0.615<br>17.250 | <br>0.002<br>0.633<br>13.880 |

[a] Highest resolution shell.
